# Utilization of a Clark electrode device as a respirometer for small insects: a convincing test on ants allowing to detect discontinuous respiration

**DOI:** 10.1101/2023.10.31.564912

**Authors:** Maïly Kervella, Céline Cansell, François Criscuolo, Frederic Bouillaud

## Abstract

Respirometry provides a direct measure of an organism’s respiration, which is a significant component of its metabolic rate. Amongst ants, variations in lifespan between different social castes (such as workers and queens) can be substantial, varying according to the species. As metabolic rate is higher in short-living species, we aimed to establish how metabolic rate and longevity may have coevolved within ant’s casts. As a first methodological step, we validate here the use of a Clark electrode initially design for measuring mitochondrial respiration control pathways, for flow-through oxygen consumption in ant, by comparison with stop flow oxygen consumption and carbon dioxide production utilizing the indirect calorimetry methodology. The global aim is to provide a reliable methodology to conduct accurate comparisons of metabolic rates within and among ant species. As expected, using Clarck electrode entails high time resolution and revealed that queens and workers exhibited discontinuous respiration, with episodes of apnea up to 20 minutes.

## Introduction

Respirometry is a technique used to measure the metabolic rate in a wide range of organisms. Aerobic respiration is a fundamental process by which these living organisms break down organic molecules (carbohydrates, peptides and lipids) to produce energy, primarily in the form of adenosine triphosphate (ATP). The rate at which an organism utilizes oxygen to produce energy through aerobic respiration to sustain their vital functions and living activities is called metabolic rate [1]. Depending on the recording conditions and physiological status of the individuals, metabolic rate can be measured as BMR for Basal Metabolic Rate, as RMR for Resting Metabolic Rate, as FMR for Field Metabolic rate, or as SMR for Standard Metabolic Rate primarily for invertebrates and poikilothermic vertebrates [2]. In brief, metabolic rate can vary between individuals, species, and even different physiological states under circadian cycle. For example, a resting animal will have a lower metabolic rate compared to the same animal when it is active. Metabolic rate can be measured by different methodologies [3]. First, it can be assessed by direct calorimetry, *i*.*e*. the measure of the amount of energy expended in the form of heat generated by the body in a calorimeter. Unfortunately, it requires very expensive equipment and may not be used in field studies [4]. With indirect calorimetry (thereafter called respirometry [5]), a measure of the heat generated by the organism is obtained via measurements of O_2_ consumption and/or CO_2_ production associated with the oxidation of energy substrates by the organisms. Gas measurements have the advantage of providing the nature of the substrates that are oxidized when the two gas are simultaneously quantified, allowing the calculation of the respiratory quotient, or RQ, the ratio of CO2 produced to O2 consumed (which is the highest for carbohydrate substrates). Indirect calorimetry also informs on metabolic efficiency, which is the number of ATP produced per O_2_ consumed. Metabolic efficiency depends not only on the type of substrate oxidized, but also on metabolic state or exercise intensity, both implying additional anaerobic pathways that increase the RQ (CO_2_ increase without O_2_ consumption [6–8]). Thus, respirometry serves as a valuable tool to assess the variations and the efficiency with which individuals modulate their metabolic activities and energy requirements, in response to environmental challenges. Because those regulations are likely to be affected by life stages (*e*.*g*., reproduction) and age, respirometry has proven itself to be a key methodology when focusing on organisms’ health and fitness (*i*.*e*., individual relative reproductive success over life [9,10]). Getting metabolic data is then crucial for understanding various ecological [11,12], physiological and evolutionary processes [10], having applications in numerous fields from functional ecology up to medical research, for instance in the study of respiratory diseases in humans [13]. Most past studies used the measurement of CO_2_ release rather than O_2_ consumption for small animals [14]. This is because the lower concentration of CO_2_ in the atmosphere (0.4%) makes it easier to detect small changes, allowing for more precise measurements during respirometry experiments and reduced interferences (CO_2_ sensors are less sensitive to humidity). Furthermore, CO_2_ is a more accurate proxy of the metabolic processes because it reflects the complete oxidation of substrates, and gives a comprehensive picture of the total metabolic activity and energy expenditure of an organism. On the other hand, O_2_ consumption is a direct indicator of an organism’s aerobic metabolic rate. In the case of studying oxygen consumption and aerobic metabolic rates, an oxygen analyser is thus more appropriate. Still, respirometry can be challenging to detect accurately gas exchanges in small organisms due to small volume of respiratory gases. One solution is to pool individuals. Additionally, the stress induced during the measurement process can further affect the reliability of the results. Social insect colonies have genetically closely related individuals that however may exhibit significant differences in size, behaviour, physiology, and life expectancy. For example, social insects’ queens can live up to 10-30 times longer than workers in some species, 800 times for extreme cases [15,16]. Among social insects’, ants are therefore a relatable model for studying ageing mechanisms while avoiding the confounding factors of different genetic background [15]. Interspecific comparisons of metabolic rate and longevity covariation has yielded the general observation that metabolic rate is higher in short-living species [17]. However, there are very few existing studies assessing how the metabolic rate and longevity may have coevolved within a given ant species, *i*.*e*. among individuals of different social casts assuming distinct social roles in the colony. In this respect, studies addressing different proxies of metabolic differential regulations according to the social role have been recently published [18–21]. Proteins and metabolites associated with metabolic activity are for instance more abundant in *Lasius niger* workers[18,20], which is in line with a higher metabolic rate measured in some species [21], raising the question of the evolution of a specific trade-off between energy metabolism and longevity in eusocial species, beyond the classical predictions of the free-radical theory of ageing [22] (*i*.*e*., notably the ability to minimize the cost of reproduction [23]).

Our laboratory conducted a project on the comparative study of ageing mechanisms among social casts of insects, based on a measure of metabolic rate in different ants’ species. As mitochondrial respiration was assessed using a Clark electrode system (Oxygraph O2k, Oroboros Instrument GmbH, Innsbruck, Austria), the same system was tested for indirect calorimetry. A Clark electrode is a polarographic oxygen sensor usually used for measuring the concentration of dissolved oxygen (DO) in a liquid. It was developed by Leland C. Clark Jr. in the 1950s and has since become a widely used tool in various scientific and industrial applications such as water treatments and biomedical research [24,25]. In 1968, Hayes and collaborators used modified Clark-type electrodes to measure the O_2_ consumption in the gaseous phase for insects weighing between 50 and 500 mg. The results were of the same order of magnitude as those obtained with the manometric system [26]. Oroboros instrument presented Figures 1 (pictures) and Figure 2 (Clark electrode functioning) are used to assess mitochondrial and cell bioenergetics through OXPHOS (mitochondrial oxidative phosphorylation system) thanks to accurate and real-time measurements of DO. The electrode consists of a cathode and an anode separated by an electrolyte, a potassium chloride (KCl) solution. The cathode is made of platinum/gold, while the anode is silver. The electrolyte allows the movement of ions between the two electrodes. When a voltage of 600 to 800 mV (potential for reducing O_2_) is applied to the Clark electrode immersed in a liquid sample containing dissolved O_2_, O_2_ molecules diffuse through a gas-permeable membrane, and reach the cathode surface where they undergo a reduction reaction. This results in an electrical current whose intensity is proportional to the concentration of dissolved oxygen in the sample (or oxygen pressure). By measuring this current, which is calibrated from 0% to 100% oxygen, the oxygen concentration can be determined : the output of the Clark electrode is connected to a computer with the data acquisition system that converts the current into an oxygen concentration reading, taking account pressure in the chamber, temperature and pH of the solution [27]. In this study, we describe the protocol used to adapt the Oroboros instrument for measuring oxygen consumption in living ants (pools of a few mg), and compared it with currently used indirect calorimetry methods for insects, and literature.

**Figure 1:**
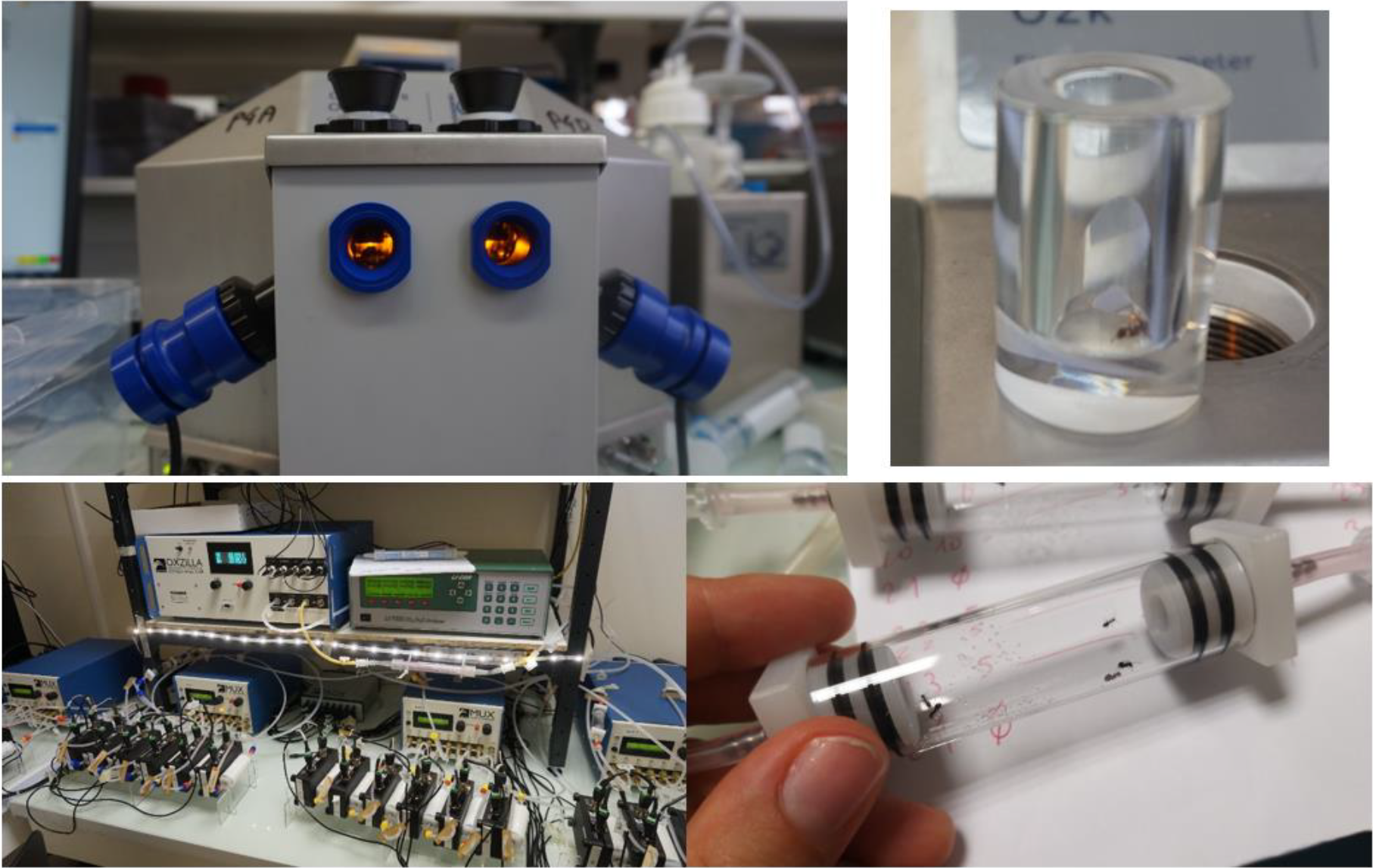
Presentation of the two different devices. Top: Oroboros with two acquisition O2k-sV chamber (0.8 cm^3^) and (right) anO2k-sV chamber containing an ant. Down: Calofly system from SABLE, with 32 chambers of 30cm^3^ (right).

**Figure 2:**
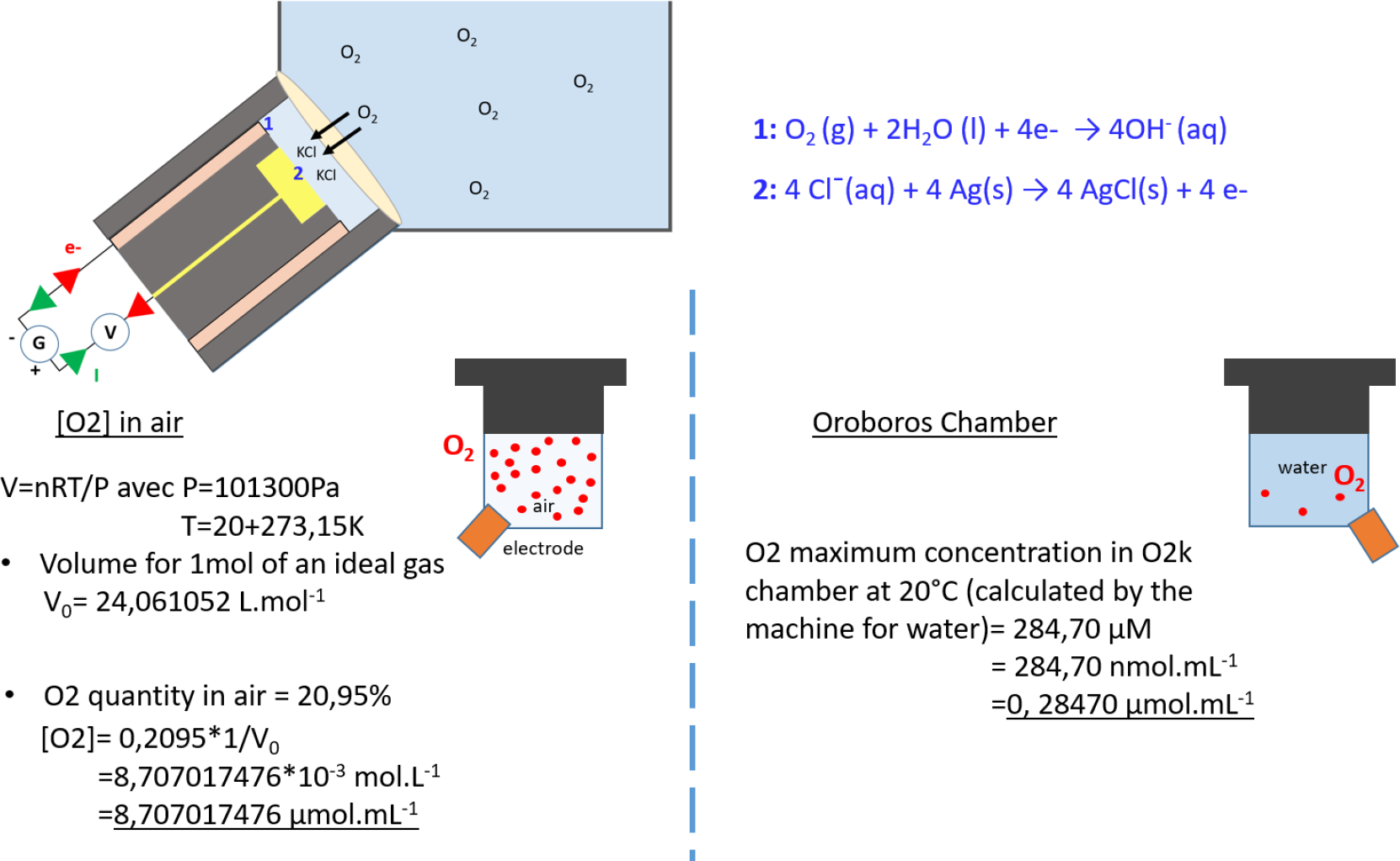
Top: Functioning of the Clark electrode. Dissolved oxygen in water pass into KCl solution via a Teflon membrane. Under the influence of the redox potential, the dissolved oxygen is ionized into hydroxide at the cathode (in pink) upon contact; Cl-precipitate at the anode (in yellow) to form AgCl. As a result, a very weak current is generated, proportional to the concentration of oxygen in the environment surrounding the electrode. This current is then amplified and used to calculate oxygen concentration in real-time. Down: schematic conversion of units between DatLab software and samples measured with air (on the left) or liquid (on the right) in the chamber.

## Materials and Methods

Three ant species has been used during the tests: the black garden ant (*Lasius niger*, Linnaeus 1758), the carpenter ant (*Camponotus herculeanus*, Linnaeus, 1758) and red-barbed ant (*Formica rufibarbis*, Linnaeus, 1761). Colonies were maintained at room temperature (20-25°C) and natural photoperiod, and were fed *ad libitum* with sugar water and mealworms. They were fasted the day before measurements. Workers were randomly chosen, *i*.*e*. mixing foragers and nest-workers. For *Lasius niger*, 1 to maximum 13 individuals were pooled per chamber. This size of pooled sample was chosen while taking into account the diameter of a chamber and the fact that *Formicidae* can release formic acid when they are stressed. In comparison, queens were placed alone, as well as workers and queens *of Camponotus herculeanus* and *Formica rufibarbis*.

### Oroboros oxygen consumption set-up

Ant’s O_2_ consumption was measured using an Oxygraph O2k (Oroboros Instrument GmbH, Innsbruck, Austria) and DatLab software. The system is usually used with stirred liquid suspensions such as isolated mitochondria, and calibrated with water solution equilibrated with ambient air oxygen (100%) and by addition of sodium disulfite in excess, a chemical consuming all oxygen present, for the 0% oxygen reference point. In this case, to proceed with a gaseous atmosphere, the electrodes were calibrated with empty chambers open to ambient air (100%) or closed and filled with argon for 0%. We therefore assumed that air saturated with oxygen (20.95% O_2_ in air) results a signal equivalent to that when the electrode is in contact with a 100% saturated water (270μmol.L-^1^), expressed in the O2k software with units in pmol.mL^-1^. The calculation is mapped Figure 2. The molar volume of an ideal gas is V_0_= 24,22 L.mol^-1^ at 20°C (293,15K) and 101,3kPa. Taking account that O_2_ quantity in air is about 20,95%, [O_2_]= 0,2095*1/V_0_ =8,65 μmol.mL^-1^, and the conversion factor between O2k units is (8.65 μmol.mL^-1^) / (0.270 μmol.mL^-1^)= 30,58. Oxygen flux obtained with the machine were considered without the background correction automatically proposed by DatLab, that correct instrumental effects: 1) the electrode oxygen consumption by itself and 2) oxygen back diffusion in a liquid [28]. Indeed, oxygen consumption by the Clark oxygen electrode has fewer consequences in air than in water due to the 30,58 times higher oxygen content in air. In addition, back diffusion was neglected because of sealing of the chambers with agarose (chambers are not normally completely sealed since stoppers have a section allowing injections with Hamilton syringe). Empty chambers were sealed at the end of each run days, and oxygen consumption by the polarographic sensor itself remained quite constant as shown Figure 4.

### Ant oxygen consumption session

We proceeded as follow: ants were isolated from their colony in a tube, temporary anesthetized on ice (less than a minute to allow an easier manipulation of individuals) and placed in a small sized respiration chamber (v = 0.800 mL, 12mm Ø). Chambers were completely sealed with agarose 2.5% casted in stoppers. Real-time O_2_ consumption was recorded at 20°C for 40 min after agarose polymerization, and respiration rate was obtained as mean O_2_ consumption during all 40 minutes acquisition (total time in the chamber about an hour).

### Carbon dioxide production on Sable device

Ants’ metabolism was assessed using indirect calorimetry with the Calofly device (as shown in pictures Fig. 1), which allowed for simultaneous measurement of metabolic activity (Sable Systems Europe GmbH). In brief, the setup consists of 32 respirometer chambers of 30 cm^3^ connected to flow devices for generating the main flow (37mL.min^-1^). Li CO2 analyzer (Li-COR Biosciences) and an Oxzilla O2 analyzer (Sable Systems) were used. The flow was calibrated using an O_2_ and CO_2_ mixture with known concentrations (Air Liquide, S.A. France). The device was operated in a stop-flow mode with a 128-minute cycle analysis, under constant darkness (due to activity measurements) and at room temperature (26°C). Values were regularly compared to a reference empty chamber. Dedicated software ExpeData was used to obtain final metabolic values.

## Results / Discussion

### The Clark electrode provides suitable results

In Figure 4A, an example of an acquisition run is presented, demonstrating the mean oxygen consumption of a pool of 6 *Lasius niger* workers as 2.8956 pmoL.s^-1^.mL^-1^ on the software (equivalent to 6.2996 μL.h^-1^ after conversion, either 1.050 μL.h^-1^ per individual). The layout for the empty control cell is shown in Figure 4B, indicating a zero consumption rate of 0.0945 (0.21 μL.h^-1^). Our experimental sample size was driven by the need to avoid social stress effects, considering that social isolation can induce stress, potentially leading to an increase in metabolic rate. For instance, using a higher sample size of workers can potentially lead to stress and formic acid spread, which may interfere with gas measurements (data not shown). To investigate the influence of the device used on the O_2_ flux, a mixed linear model was run (*lmer* function from *lme4* R package) for the three ant species including workers and queens (66 samples measured with Oroboros device, 37 with the Sable device). Results are presented in Figure 3A. Power laws were fitted on log-transformed data of O_2_ consumption flux as the response variable, and log(pool body mass) and device as factor variables, including interactions term. Because ant groups are heterogeneous, species and caste were used as a random factors. The *anova (type III)* function was then performed on the model to obtain the p-value of each explanatory variable. Parameter estimates and associated standard errors are given Figure 3B. The main conclusions are that O_2_ consumption scaled to body mass as expected [29,30], and that there is no statistical difference between the two machines used. However, the Clark electrode values seems to be slightly higher for lower body mass, and lower for the higher body mass (interaction term, p = 0.096). Metabolic measurements were not all done at a same temperature (temperature is not controlled in the Sable device), and thus the temperature effect is a confounding factor with the device. A temperature effect may have driven part of the differential metabolic results we get with the Sable device, since increased ambient temperature is known to impact ectotherms’ physiological traits [29,31,32]). Still, our data suggests that, if there was a temperature effect on metabolic measurements, this remained non-significant. Figure 3C depicts the consistent mean oxygen consumption per *Lasius niger* individual across a range of group sizes, from 1 to 13 individuals. Comparing the three sample sizes with at least four biological replicates (n=5-7, Fig. 3C) led to no significant differences in oxygen consumption with the number of ants present in the chamber (F=1.809, dF=2, P value=0.184). The mean oxygen consumption per individual, measured using the Clark electrode was 0.9851 ± 0.2325 μL.h^-1^. When a single individual was introduced, the measured oxygen consumption appeared to be doubled (Fig. 3C). This may reflects the stress effect linked to isolation as ants are social insects. An alternative explanation for this rise in oxygen consumption could be that social isolation correspond to a foraging situation in ants, which may contribute to an increase in metabolism among more active foragers. Unfortunately, locomotor activity was not assessed in the Oroboros. Most data were obtained for the sample size of 6 individuals. Values for these 21 replicates were heterogenous, with a maximal variation of 2,5 for oxygen consumption. It is not surprising to have variations between individuals: they are not of same age, do not have same task in the nest even among workers (there is an age-related division of labour in the species), and short-term variability in body mass may affect our measurements, as metabolic rate was found to covary with individual mass. Furthermore, body composition changes with age and caste [33]. In litterature, Jensen used in the 70’s the Warburg technique (based on pressure change in a closed system) to measure ant oxygen consumption [34]. They found individual *Lasius niger* consumme 1.08 ± 0.50 μL.h^-1^ at room temperature, and 0.68 μL.h^-1^ at 20°C [35] (see Table 1). *Lasius niger* workers had a similar O_2_ consumption between Oroboros and the Warburg system at RT, while being 19.4 % higher than with the gas analyser Sable. At 20°C, the Oroboros gave 44% higher O_2_ consumption than the Warburg apparatus. Queens *Lasius niger* had an oxygen consumption 27% higher using the Oroboros than using the gas analyser. Results between the two respirometers tested here were comparable for *Formica rufibarbis* workers. Oxygen consumption values obtained using the Oroboros were lowered by a factor two for *F. rufibarbis* queens, and even by three for the *Camponotus herculeanus* queens. Prolonged periods of apnea were observed in the latter case, coresponding to the pattern of discontinuous gas exchange (DGC) as shown in Figure 4. Periods between two burst of oxygen consumption can encompass half of the acquisition time and this can partly influence regression lines. Whether this may also account for differences in metabolic measurements between devices needs further study.

**Figure 3:**
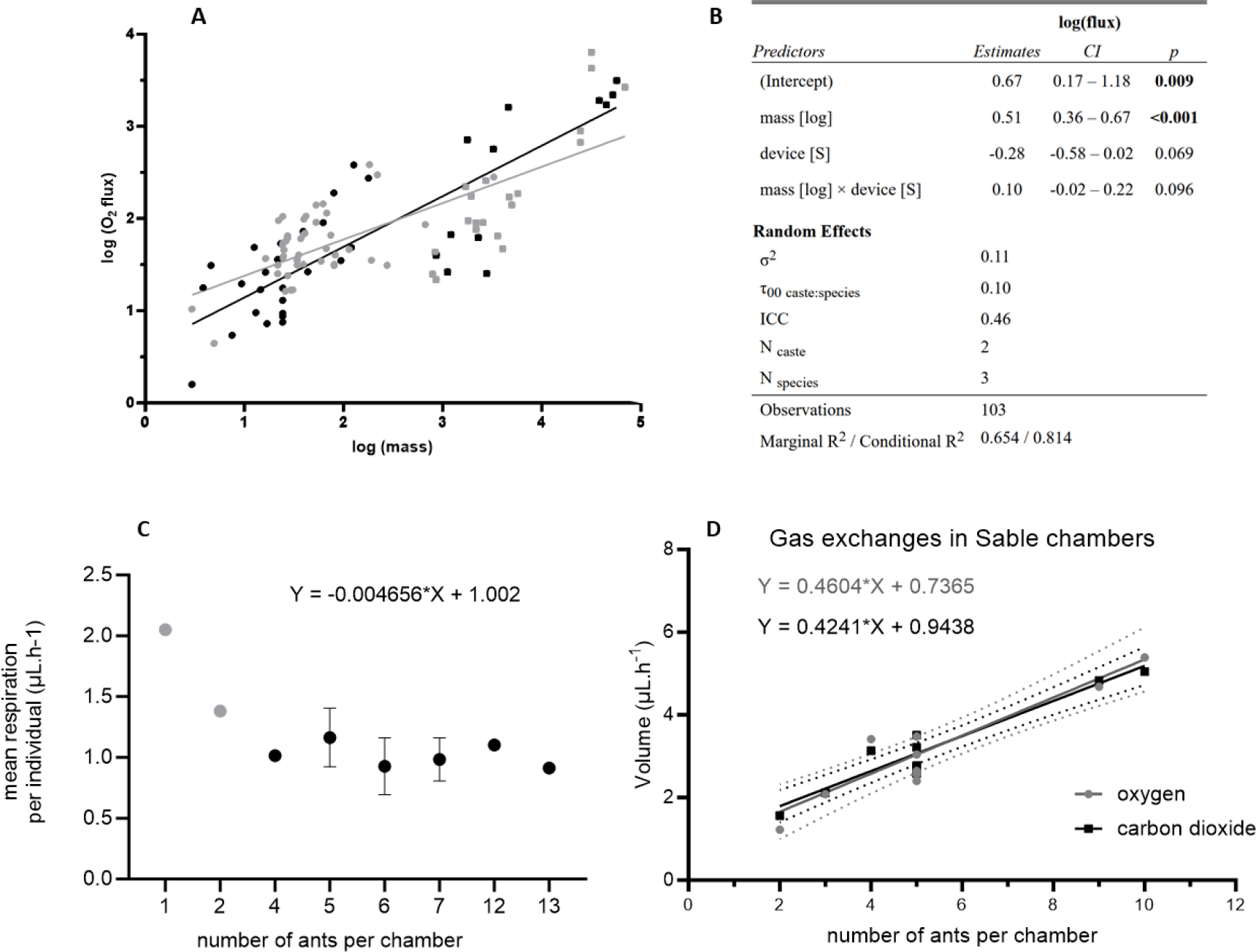
A: Comparison of oxygen consumption measurements obtained using the Clark electrode (Oroboros, 20°C acquisition, in black) and the Sable system (room temperature, in grey). Circles represent ant workers, squares queens. B: Parameter estimates and associated standard errors of the Mixed Linear Model used to test for differences in metabolic measurements between the two devices. C: Mean oxygen consumption *per* ant in Oroboros chambers depending on the number of individuals (mean ±SD). D: SABLE measurements, where O_2_ and CO_2_ were measured concomitantly to determine RQ in pools of *Lasius niger* in relation to sample size of ants present in the chambers.

**Figure 4:**
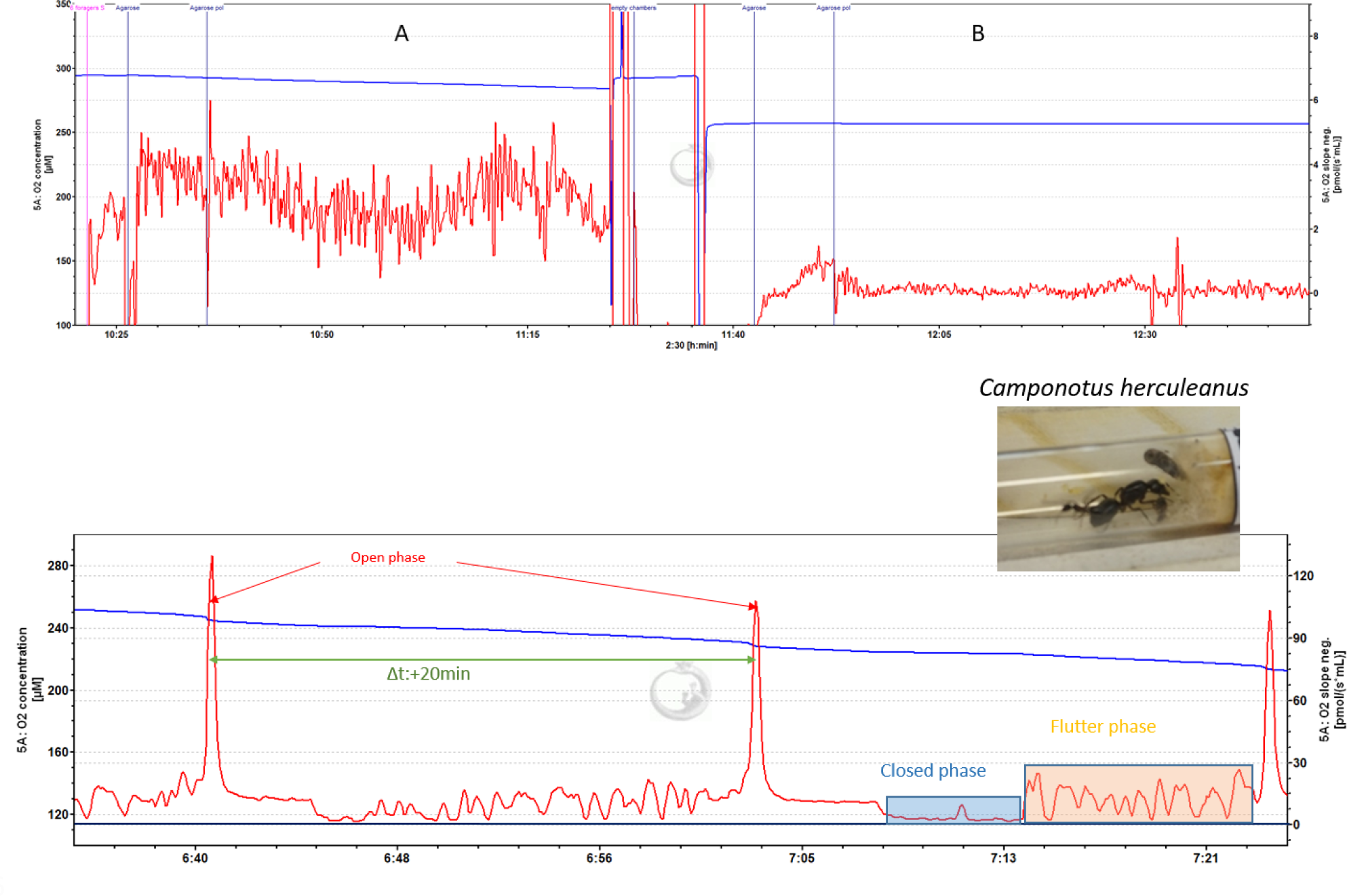
Top: example of a session recording respiration of 6 workers of *Lasius niger* on DatLab software (A), followed by empty chambers sealed to verify electrode derivation (B). Down: surprisingly, the Clark electrode permited to observe discontinuous gaz exchanges in some queens and workers from the biggest individuals (i.e., the oxygen pic during open respiration phase). Here, an example on a *Camponotus herculeanus* queen is presented, for which all different phases of respiration already described in insects were observed (open phase with pic of oxygen, closed phase when spiracles do not permit gas exchanges at all, flutter phase with fast fluctuating O_2_ consumption).

**Table 1:** Gas exchanges measurements in three *Formicinae* species, in workers and queens. Comparison between the two devices tested, and values found in *Jensen et al*., 1975 in grey [34] or Jensen 1978 [35]. * indicates when the device measured O_2_ consumption, but non-reliable ants RQ.

| Species | Caste | Material | O<sub>2</sub> | CO<sub>2</sub> | Respirometry method | T°C | n | mean | SD |
| --- | --- | --- | --- | --- | --- | --- | --- | --- | --- |
| *L. niger* | W | Clark electrode | ✓ |  | flow-through | 20 | 33 | 0.9851 | 0.2325 |
| *L. niger* | W | gas analyser | ✓* | ✓ | stop-flow | RT | 15 | 0.8251 | 0.4095 |
| *L. niger* | W | manometer | ✓ |  | stop-flow | RT | RT | 1.08 | 0.58 |
| *L. niger* | W | manometer | ✓ |  | stop-flow | 20 | RT | 0.68 | ND |
| *L. niger* | Q | Clark electrode | ✓ |  | flow-through | 20 | 8 | 6.147 | 2.138 |
| *L. niger* | Q | gas analyser | ✓* | ✓ | stop-flow | RT | 4 | 4.846 | 0.9922 |
| *C. herculeanus* | W | Clark electrode | ✓ |  | flow-through | 20 | 7 | 4.989 | 0.9015 |
| *C. herculeanus* | W | gas analyser | ✓* | ✓ | stop-flow | 26 | 5 | 9.579 | 2.867 |
| *C. herculeanus* | W | manometer | ✓ |  | stop-flow | RT | RT | 13.56 | 5.57 |
| *C. herculeanus* | W | manometer | ✓ |  | stop-flow | 20 | RT | 7.90 | ND |
| *C. herculeanus* | Q | Clark electrode | ✓ |  | flow-through | 20 | 6 | 30.34 | 10.75 |
| *C. herculeanus* | Q | gas analyser | ✓* | ✓ | stop-flow | 26 | 4 | 107.6 | 8.248 |
| *F. rufibarbis* | W | Clark electrode | ✓ |  | flow-through | 20 | 5 | 4.183 | 2.019 |
| *F. rufibarbis* | W | gas analyser | ✓ | ✓ | stop-flow | 26 | 5 | 4.157 | 1.132 |
| *F. rufibarbis* | Q | Clark electrode | ✓ |  | flow-through | 20 | 6 | 9.958 | 1.163 |
| *F. rufibarbis* | Q | gas analyser | ✓* | ✓ | stop-flow | 26 | 4 | 15.94 | 7.682 |

### Calculation of RQ and Circlesrcadian cycle effects on Sable device

Similar slopes were obtained for O_2_ and CO_2_ flux for different pool sizes of *Lasius niger* workers (Figure 3B). Since oxygen consumption is very low in such models, ΔO_2_ after chamber opened do not decrease markedly and can be slightly misestimated compared to ΔCO_2_. Therefore, most RQ obtained were higher than 1 (corresponding to lipid synthesis), with no evolution during time despite ants were fasting into the chambers during more than 24 hours (see Table 2). Hence it was not possible to determine the nature of substrate oxidized. In other insects like flies (note that metabolic rate is higher than in ants [36,37]), while CO_2_ can be assessed accuratly in a single fly, flies were pooled *per* 25 individuals to measure O_2_ more accuratly in order to determine RQ [38].

**Table 2:** RQ obtained during the Sable system acquisition. The cases en grey correspond to RQ>1. n is the number of ants per chamber. Numbers indicated after RQ correspond to the chamber ID (see also Figure 6).

| n | 5 | 10 | 5 | 5 | 5 | 2 | 5 |
| --- | --- | --- | --- | --- | --- | --- | --- |
|  | RQ_18 | RQ_19 | RQ_22 | RQ_26 | RQ_27 | RQ_28 | RQ_31 |
| 16:08:04 | 0,9843771 | 0,8479304 | 1,05812 | 1,139547 | 1,009154 | 1,434737 | 1,076952 |
| 18:16:04 | 1,035072 | 0,8973882 | 1,004417 | 1,040707 | 0,9057006 | 1,372361 | 0,9867143 |
| 20:24:04 | 0,9778897 | 0,8655667 | 0,9245234 | 1,166726 | 0,9262109 | 1,133282 | 0,9705676 |
| 22:32:04 | 0,7252429 | 0,9371588 | 0,8793747 | 1,164221 | 0,8486915 | 1,312386 | 0,8047348 |
| 00:40:04 | 1,033311 | 0,8800876 | 0,7044209 | 1,266591 | 0,9256601 | 1,529205 | 0,8965204 |
| 02:48:04 | 1,007793 | 0,9459545 | 0,9002062 | 1,567793 | 0,9969299 | 1,206778 | 0,8154089 |
| 04:56:04 | 1,009547 | 0,9583928 | 1,020491 | 1,234648 | 1,027962 | 1,039096 | 1,064386 |
| 07:04:04 | 0,9479125 | 0,8497361 | 0,8983144 | 1,326314 | 1,137401 | 1,263359 | 0,9920726 |
| 09:12:04 | 1,074703 | 0,9157327 | 0,9480888 | 1,36398 | 1,071026 | 1,230585 | 1,066416 |
| 11:20:04 | 1,020342 | 0,8937598 | 1,012466 | 1,226922 | 1,021975 | 1,243248 | 0,9987755 |
| 13:28:04 | 0,9194415 | 0,9595323 | 0,8464941 | 1,261465 | 0,9967823 | 0,9980044 | 0,9851054 |
| 15:36:04 | 0,9130132 | 0,895824 | 0,9390872 | 1,362157 | 1,005835 | 1,290588 | 23,2103 |
| 17:44:04 | 1,07469 | 1,102292 | 0,8578913 | 1,125283 | 0,9750324 | 0,9253665 | 0,9649587 |
| 19:52:04 | 0,9607995 | 0,9448576 | 0,8842281 | 1,22971 | 0,9579208 | 1,136717 | 1,143873 |
| 22:00:04 | 1,107666 | 0,9464912 | 0,8810239 | 1,305277 | 1,038764 | 1,063407 | 1,132444 |
| 00:08:04 | 1,021079 | 0,9591442 | 0,8386973 | 1,334618 | 1,143914 | 1,870823 | 1,372764 |
| 02:16:04 | 0,9762473 | 1,035334 | 0,9532833 | 1,414549 | 1,145106 | 1,66622 | 0,921229 |
| 04:24:04 | 1,11228 | 0,9954199 | 0,8597758 | 1,254105 | 1,086647 | 2,125256 | 1,155826 |
| 06:32:04 | 5,791422 | 0,9693155 | 0,963133 | 1,323314 | 1,028135 | 1,638574 | 1,238518 |

To assess any circadian rhythm in metabolic rate, a Kwiatkowski-Phillips-Schmidt-Shin (KPSS) was run on O_2_ as CO_2_ repeated measurements conducted over 36h, using the *tserie* R package Respiration rate (both using O_2_ as CO_2_ values) was significantly constant during the 36h acquisition, as shown in Figure 5, and no circadian rhythm of respiratory metabolism was observed. It can be due to the fact that heterogenous workers were merged (age and behavior), excluding the possibility to see a social synchronization during the 36 hours acquisition, as it has been shown in honeybee [39]. Note that using our data on activity behaviour (Figure 6), we do not show neither rhythmicity, which can be surprising based on past studies in other ants. For instance, *Camponotus rufipes* foragers showed circadian foraging behaviour synchronized with food availability [40]. Japanese *Diacamma indicum* also exhibit circadian activity rhythms, showing typical actograms that depend on the tasks within the colony. In the present study, there were no food available, no pupae to take care of, no queen present *i*.*e*., all being factors that can act as social rhythm regulators. In addition, young workers are found to be twice as inactive as older ones, displaying a more pronounced daily rhythm [41]. Rhythms also depend of the brood type being taken care of [42]. Interestingly, workers that interacted with different-aged individuals showed a reduction of circadian rhythmicity [43]. A mixed linear model was employed, with CO_2_ flux as the response variable (as it is more sensitive than O_2_ levels). Both the activity level and the number of ants per chamber were introduced as fixed factors. However, activity level did not exert a significant influence on CO_2_ levels. Overall, our results on O_2_ as CO_2_ measurements are not supporting the hypothesis that there is a circadian rhythm in metabolic rate in our species, or that increased levels of locomotor activity lead to higher oxygen consumption.

**Figure 5:**
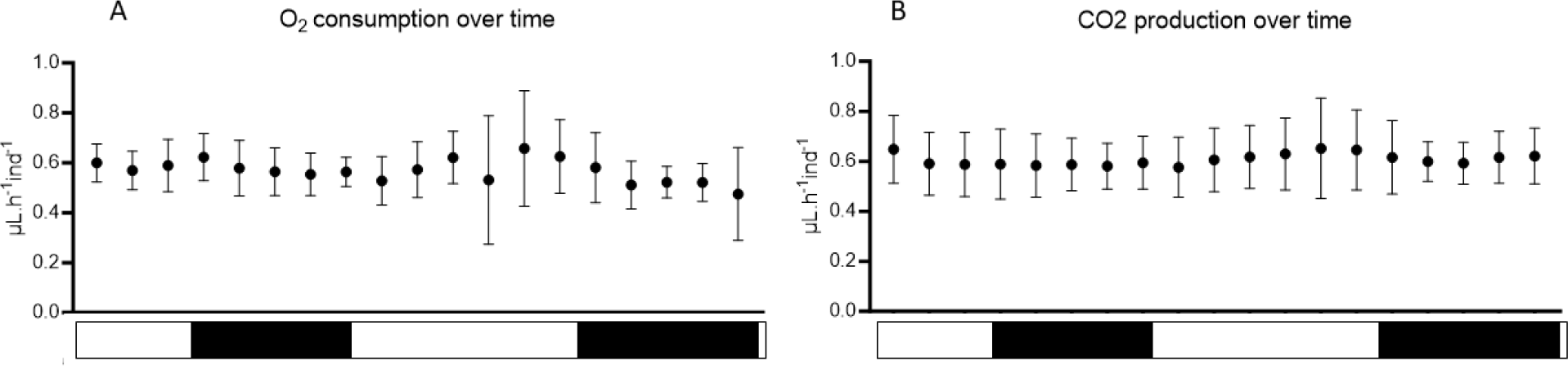
mean O_2_ consumption (A)/ CO_2_ production (B) per ant during 36 hours acquisition. White bars labelled “Day”; black bars labelled “Nightime”, but ants were in continuous dark due to activity detectors. Measurements have been conducted on 7 chambers with pool of 2 to 10 individuals.

**Figure 6:**
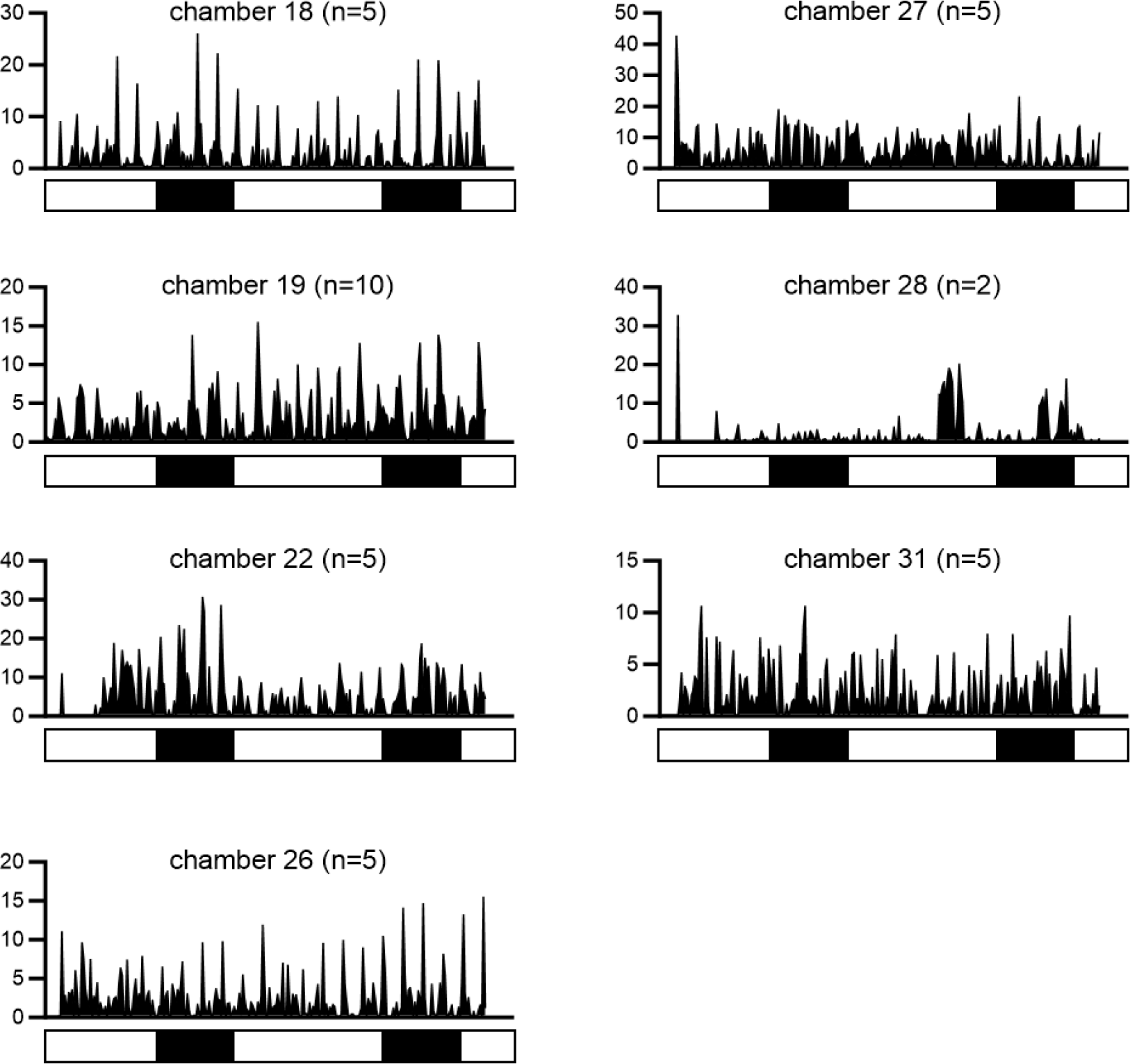
Ants’ locomotor activity measured in the seven chambers used for RQ and circadian rhythm measurements by the SABLE system (numerous of the chamber used table 2). White bars labelled “Day”; black bars labelled “Nightime”, but ants were in continuous dark due to activity detectors. Relative activity for 10 min is given in ordinate.

### The Clark electrode allow to analyse discontinuous gas exchanges in larger species

Interestingly, the flow-through system allowed to observe discontinuous gas exchanges (DGC, Fig. 4) in the biggest ants (*Camponotus herculeanus*). This has been previously described in others ant models, notably *Camponotus* species [44,45]. With the system being closed and lacking any gas homogenization, as achieved with a stirrer in an aqueous phase, the Clark electrode gave distinct and clear bursts of O_2_ (slightly smoothed), making our protocol highly adapted to the study of discontinuous respiration in insects. There are a few comparisons of discontinuous respiration among castes of a same species in literature: within the same colony, ant castes may exhibit different gas patterns depending on the habitat characteristics and caste roles [46,47]. Lighton and Berrigan discuss the fact that queen ants are confined to underground and hypercapnic chambers. Worker ants, in contrast, move between these chambers and the surface for colony tasks and should then have a normoxic metabolism. As a result, queens are predicted to likely use DGC during anoxia, thereby also reducing respiratory water loss. For foraging workers, the DGC probably helps to also lower transpirational water loss rates based on external conditions [48]. Based on available evidence, the DGC could potentially serve multiple adaptive functions, although their general occurrence remains uncertain [49]. Another explanation that befits the metabolic rate and oxidative stress theories for social organisms is the oxidative damage hypothesis. The closed phase is characterized by a rapid decrease in tracheal oxygen concentration to low partial pressures, while a crucial flutter phase ensures oxygen supply matches tissue consumption rates [50]. Furthermore, metabolic rate has been shown lowest for queens using DGC in *Pogonomyrmex barbatus*, intermediate for individuals using cyclic gas exchange, and highest for individuals using continuous gas exchange [51]. Consequently, DGC may lower oxidative stress. Matthews and White also suggested that queen ants might also exhibit DGCs due to reduced brain size and activity related to their limited behavioral repertoires and a lack of visual stimulation found in their claustral condition [52,53].

### General conclusion

The study confirmed the Clark electrode’s suitability for quantifying oxygen consumption in ants, successfully capturing DGC in larger ants. This device offers temperature control and quasi-continuous measurements (1 measurement every 2 seconds for the shortest interval). However, it only provides two measures per run, and due to sealed chambers, additional waiting time is needed for agarose polymerization, which can be time-consuming for numerous insects. Moreover, while the results seem promising, it raises the question of whether they are sufficient to detect differences among workers with distinct social roles. For investigating DGC in smaller individuals, it implies that they are placed alone in respirometry chambers, and we highlighted putative stress effects on gas consumption. In that case, a more sensitive alternative system might be preferable, with a controlled flux allowing to describe in details DGC phases are very short in time.

## Authors’ contributions

MK and CC did measurements, MK and FC wrote a first version of the text and all co-authors drafted the final manuscript and gave their approval for publication.

## Acknowledgements

We acknowledge the technical platform Functional and Physiological Exploration platform (FPE) of the Université Paris Cité, CNRS, Unité de Biologie Fonctionnelle et Adaptative, F-75013 Paris for generously providing us access to their Sable device.

